# Stratified Test Accurately Identifies Differentially Expressed Genes Under Batch Effects in Single-Cell Data

**DOI:** 10.1101/2021.06.08.447617

**Authors:** Shaoheng Liang, Qingnan Liang, Rui Chen, Ken Chen

## Abstract

Analyzing single-cell sequencing data from large cohorts is challenging. Discrepancies across experiments and differences among participants often lead to omissions and false discoveries in differentially expressed genes. We find that the Van Elteren test, a stratified version of the widely used Wilcoxon rank-sum test, elegantly mitigates the problem. We also modified the common language effect size to supplement this test, further improving its utility. On both simulated and real patient data we show the ability of Van Elteren test to control for false positives and false negatives. A comprehensive assessment using receiver operating characteristic (ROC) curve shows that Van Elteren test achieves higher sensitivity and specificity on simulated datasets, compared with nine state-of-the-art differential expression analysis methods. The effect size also estimates the differences between cell types more accurately.

## 1 Introduction

Large-scale studies such as the Human Cell Atlas [1] involve hundreds of laboratories, thousands of patients, and millions of cells, bringing about both opportunities and challenges in analyses. When comparing cell types or groups, discrepancies across experiments and differences among participants lead to omissions and false discoveries in differentially expressed genes. Even the trend (upregulated or downregulated) can be reversed in a phenomenon called Simpson’s paradox [2]. These phenomena are termed “batch effects”. For differential expression analysis, batch effects can be handled before performing statistical testing, or factored into the test. Although multiple methods have been proposed to tackle the batch effects, no such option for the widely used Wilcoxon rank-sum test [3], [4] has been applied to single-cell studies, to the best of our knowledge. Here, we show that the stratified rank-sum test (known as Van Elteren test [5]) and our modified common language effect size may fill this gap and benefit single-cell studies.

We briefly review and conceptually compare related works on correcting batch effect in section 1.1. Then, in section 2, we revisit Wilcoxon rank-sum test, and introduce the Van Elteren test supplemented by our direct extension of the common language effect size [6], [7]. In section 3, we use a few examples to illustrate the scenarios where a stratified test is necessary. A comprehensive assessment on simulated datasets shows that Van Elteren test identifies differentially expressed genes with higher sensitivity and specificity, compared with nine state-of-the-art differential expression analysis methods. We also show an application to retinal data. The results show that controlling for batch effects in the Wilcoxon test and its corresponding effect size leads to more accurate biological discoveries, which is the major contribution of this article. More discussions and explanations are shown in section 4.

### 1.1 Related Works

Mainstream methods to mitigate batch effect fall into two categories, batch correction methods and batch-aware statistical tests. The former includes methods reducing batch effect in the data to facilitate downstream analysis, while the latter includes analyses that control for the batch effect.

#### 1.1.1 Batch Correction Methods

Batch correction methods eliminate the discrepancy among batches to create an integrated dataset. The most conspicuous manifestation of batch effect is splitting one cell type into multiple clusters. To solve this problem, many methods match and combine clusters across samples based on similarities. A commonly adopted one, Mutual nearest neighbor (MNN) [8], uses similar cells across datasets as anchors, and based on them correct the gene expression of other cells. Scanorama [9] and the integration utility in Seurat [10] are both based on the MNN methodology. Another method, Harmony [11], iteratively corrects the data by clustering the cells and moving neighboring clusters toward each other. These methods typically produce a unified data matrix, which can be conveniently used in visualization and downstream analysis. However, these empirical corrections usually lack negative control and raise uncertainty in the discovery [12]. Thus, while the corrected data help in visualization and trajectory inference, raw data is recommended for statistical testings [13]. Normalized (and log-transformed) data may also be used if necessary, but any more correction is discouraged.

#### 1.1.2 Batch-Aware Statistical Tests

Instead of manipulating the data directly, statistical methods may handle batch effect by considering it as a covariate in the model. Both classical statistical tests and models specifically designed for single-cell data have been adopted in scRNA-seq analyses. The most popular ones are integrated into Seurat [10], a widely used scRNA-seq data analysis platform. To date, it includes Wilcoxon rank-sum test, likelihood ratio test [14], ROC (Receiver operating characteristic) Analysis, Student’s t-test, negative binomial test, Poisson test, logistic regression, MAST [15], and DESeq2 [16]. Among those methods, VanElteren, Linear Regression test, Negative Binomial test, Poisson test, and MAST are capable of accounting for covariates, which can be used to model the batch effects. A more detailed description of these methods is available in Section 2.3.

Notably, all these tests are parametric, meaning that a distribution must be given in advance. However, the debate of the true distribution of single-cell gene expression has never ceased [17], which is a reason why the nonparametric Wilcoxon rank-sum test is widely used. To allow modeling covariates in nonparametric tests, one may use a generalized version of rank-sum test, the proportional odds model [18]. However, modeling batches by using a covariate also makes unnecessary assumptions upon them. Stratification, which only combines statistics from batches, is the “as simple as possible, but no simpler” way to handle batches. The Van Elteren test we use, is the stratified version of Wilcoxon rank-sum test.

It is worth noting that methods like scVI [19] have combined statistical modeling with batch effect correction. However, the effect of batches is modeled by a black-box neural network, making it subject to the same problem of batch correction methods.

## 2 Methods

### 2.1 Wilcoxon Rank-Sum Test

We briefly revisit the Wilcoxon rank-sum test (also known as Mann–Whitney U test) [3], [4]. The test statistics *U* is defined as

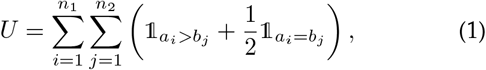

where 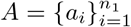 and 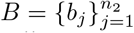 are the two samples to be compared (e.g., two cell types in one experiment), with sample sizes *n*_1_ and *n*_2_, respectively. Function 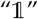 takes value 1 when its condition holds true, and 0 otherwise. When *n*_1_ and *n*_2_ are both at least 10, which is common in single-cell studies, the distribution of *U* approximately follows a normal distribution 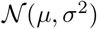 where

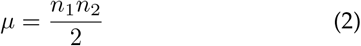

and

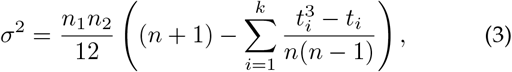

in which *t_i_* stands for ties (corresponding to the second term in Equation 1).

### 2.2 Van Elteren Test

The Van Elteren test [5] is the stratified version of Wilcoxon rank-sum test. For example, if there are *m* patients, they maybe treated as strata. In that case, a *U* statistic may be obtained from each patient *g* ∈ {1, …, *m*}, denoted as 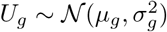. A new statistics *V* is constructed by

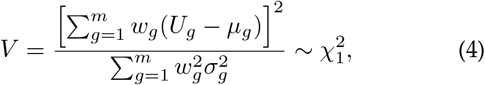

where *w_g_* is a weight for each sample to be discussed later. When *m* = 1, the formula degenerates to 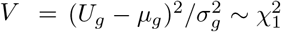, which is consistent with the rank-sum test.

#### 2.2.1 Weights

As discussed by Van Elteren [5], the weights *w_g_* can be assigned in different ways. It should be noted that the *U_g_* for a batch *g* ranges from 0 to *n*_*g*1_ *n*_*g*2_, the product of two sample sizes in the batch. Should the weights all be equal, a patient with more cells available will dominate the test results. It is proven in [5] that weight

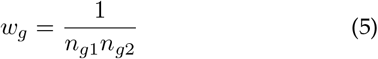

eliminates such effect, and a test utilizing such weight is thus named as “design-free test”. However, given that a batch with more instances available (e.g., a patient with more cells sequenced) may be more convincing, another weight

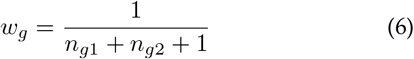

is introduced, which gives more power to larger batches. It also effectively assigns larger weights to batches whose samples are more balanced, when the batch sizes are the same. It is shown in [5] that this choice yields largest statistical power against randomized alternatives, and is thus named as “locally-best test”. The comparison of two weights are shown in Fig. 1.

**Fig. 1.**
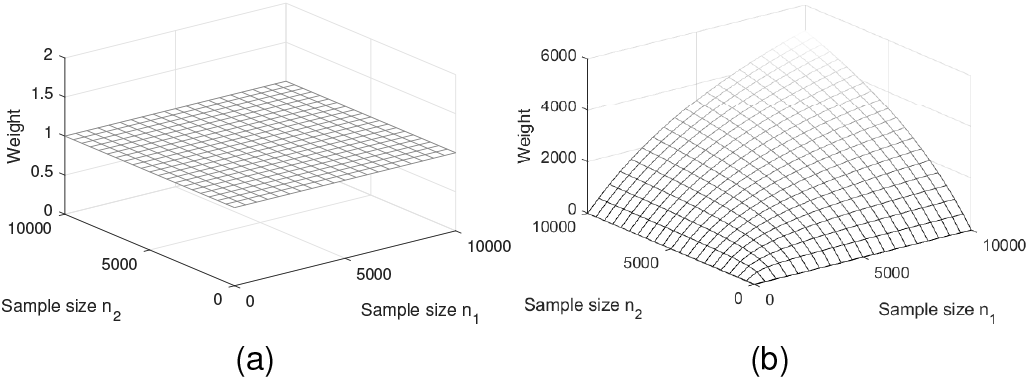
The weight (z-axis) for each batch when using the (a) design-free test and (b) locally-best test. samples sizes for two batches are on x- and y-axis. For design-free test all the batches have equal weights, while for locally-best test higher weights are given to batches with higher and balanced sample sizes.

#### 2.2.2 Effect Sizes

For Wilcoxon rank-sum test, a simple definition of effect size is

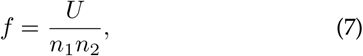

which is centered at 50%, meaning the probability *P*(*a* > *b*) when *a* and *b* are randomly drawn from sample *A* and sample *B*, respectively. An effect size greater than 50% generally means that *A* is higher, and vice versa. It may be easily extended for Van Elteren test by taking average using desired weights. For the design-free test, the effect size is

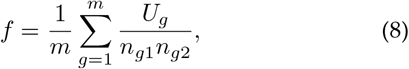

as all batches are treated equally regardless of the sample sizes. It may be interpreted as the probability of *P*(*a_g_* > *b_g_*) for *a_g_* and *b_g_* randomly drawn from *A_g_* and *B_g_*, after randomly choosing a batch *g*. For the locally-best test, the effect size becomes

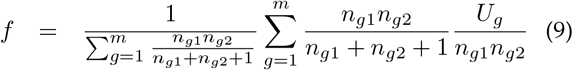

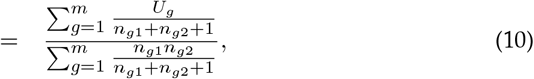

which changes the probability of choosing a group *g* to be in proportion to 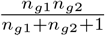, giving higher weights to batches with higher and balanced sample sizes (Fig. 1). Generally, any *w_g_* may be used to define *f*, as in

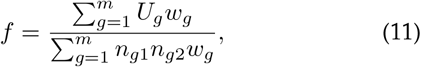

the two previous options being its special cases.

### 2.3 Other Statistical Tests

We briefly introduce likelihood ratio test [14], ROC (Receiver operating characteristic) Analysis, Student’s t-test, negative binomial test, Poisson test, logistic regression, MAST [15], and DESeq2 [16], which we compared Van Elteren test with in Section 3.2.

#### 2.3.1 Likelihood Ratio Test

The likelihood ratio test introduced by McDavid et al. [14] checks if the probabilities (*π*_1_ and *π*_2_) of cells expressed a gene and the mean expressions (*μ*_1_ and *μ*_2_) of the cells that express the gene have changed between the two groups. For each group *k* ∈ 0, 1 the likelihood is defined as

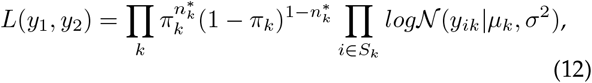

where *y_k_* is the expression profile for a specific gene, 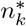 is the number of cells expressed the gene, *S_k_* is a set of the cells expressing the gene, and *σ*^2^ is the second parameter for the log normal distribution. The likelihood is maximized for the condition *H*_0_: *π*_0_ = (*π*_1_ and *μ*_0_ = *μ*_1_ and its opposite. The likelihood ratio Λ is then used as the criteria to identify the differentially expressed genes. A p-value can be derived using the fact that –2logΛ asymptotically converges to a *χ*^2^ distribution.

#### 2.3.2 ROC Analysis

A classifier is built on each gene alone to classify two groups of cells. The area under the curve (AUC) of the ROC for the classifier is used as the metric. An AUC close to 0 or 1 suggests a perfect separation, while a value of 0.5 implies a perfect mixture. The predicted “power” is defined as 2|*AUC* – 0.5| ∈ [0, 1], where higher is better. A p-value is not available for this method.

#### 2.3.3 Student’s t-Test

The distribution of the expression of a gene in both groups are assumed to follow normal distributions with unknown variances. Differentially expressed genes and their p-values are identified by Student’s t-test.

#### 2.3.4 Negative Binomial Test and Poisson Test

A negative binomial or Poisson generalized linear model (GLM) is used to identify the differentially expressed genes. Covariates may be added to account for the batches. The distribution of the coefficient accounting for two groups asymptotically converges to a t-distribution and a p-value can be derived.

#### 2.3.5 Logistic Regression

Similar to ROC analysis, a logistic regression classifier is built on each gene. A likelihood ratio test is used to calculate the p-values.

#### 2.3.6 MAST

MAST [15] is a method specifically designed for scRNA-seq data. It models the log-transformed expression. Similar to the likelihood ratio test, MAST models the number of expressed cells and the expression in the expressed cells separately. A logistic regression model and a Gaussian linear model are used, respectively. The cell detection ratio (CDR), i.e., the proportion of genes expressed in a cell, is added as a covariates to account for nuisance effects in single-cell data. Other covariates may also be manually added.

#### 2.3.7 DESeq2

DESeq2 is based on a negative binomial GLM, with more detailed modeling to address large dynamic range, outliers, etc. The p-value is derived using the Wald test.

#### 2.3.8 SigEMD

SigEMD [20] combines a data imputation approach, a logistic regression model, and a nonparametric method based on the Earth Mover’s Distance. It can also use gene interaction network information to reduce false positives.

#### 2.3.9 DEsingle

To model single-cell data, DEsingle [21] uses a zero-inflated negative binomial distribution to find genes with a significant change in the proportion of drop outs, expression levels, or both. Batch effect is not modeled.

#### 2.3.10 Accounting for Batch Effects

Poisson test, negative binomial test, MAST, and DESeq2 can account for batch effects using covariates. For MAST, we one-hot encode the categorical batches as multiple real-valued covariates, each corresponding to a batch (i.e., a patient or an experiment).

## 3 Results

We implemented the Van Elteren test with the effect size in R, available at our GitHub repository (https://github.com/KChen-lab/stratified-tests-for-seurat), based on Seurat 3.0 by utilizing its differential expression analysis part (but irrelevant to the data integration) [10]. When there are two groups of cells, A and B, denoted as type, and patient identity, denoted as batch, the Van Elteren Test can be called as follows.

~~~
FindMarkers (x, ident.1 = ‘A’, ident.2 = ‘B’, group.by = ‘type’, latent.vars = ‘batch’, test.use = ‘VE’, genre = ‘locally-best’)
~~~

The `genre` may be set to either `locally-best` or `design-free`, as introduced in section 2, based on which p-values and effect sizes are calculated. Typical results are shown in Table 2 and 3. An effect size of larger than 0.5 indicates a higher expression in cell type A, and vice versa. The `avg_logFC`, average logarithmic fold changes, are calculated automatically by Seurat, where a positive value indicates a higher expression. It may show different trends compared with the effect sizes. Generally, the effect sizes are more indicative after controlling for the batch effect.

### 3.1 Illustrations

To show the utilities of Van Elteren test, we simulated a illustrative dataset. The parameters are specified in Table 1. Poisson distribution is used to model sequencing depth. Visualization is available in Fig. 2 for illustration. We assume that the library size of each sample is equalized by other genes beyond the simulated ones. For illustrative purpose only, we applied Van Elteren test and Wilcoxon rank-sum test for an intuitive comparison, as the former is a stratified version of the latter. A more comprehensive comparison of all the state-of-the-art methods on a more realistic dataset is available in Section 3.2. The results are shown in Table 2 and 3. Trend (A over B) are indicated by arrows. For Van Elteren test, the locally-best version and the design-free version return very similar results.

**Fig. 2.**
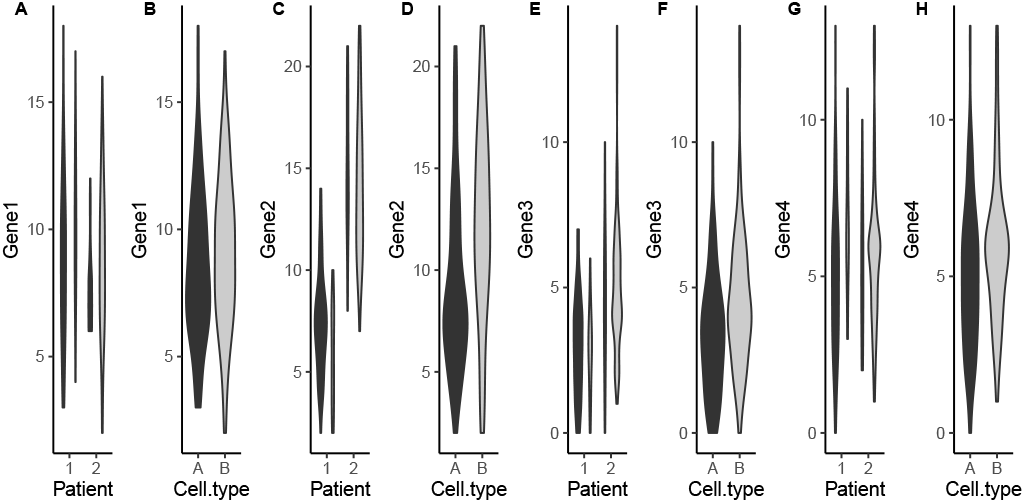
The illustrative dataset. Two shades correspond to two cell types (dark: cell type A; light: cell type B). For each gene, the left panel is stratified by patients and the right panel shows aggregated distribution.

**TABLE 1.** Rates for Poisson Distribution in Illustrative datasets

| Patient | Number 1 |  | Number 2 |  |
| --- | --- | --- | --- | --- |
| Cell type | A | B | A | B |
| Cell amount | 101 | 30 | 31 | 100 |
| Gene1 | 9 | 10 | 8 | 9 |
| Gene2 | 8 | 6 | 15 | 14 |
| Gene3 | 3 | 3 | 5 | 5 |
| Gene4 | 5 | 6 | 5 | 6 |

**TABLE 2.** P-values on the Illustrative Dataset

|  | Wilcoxon | Van Elteren |  |
| --- | --- | --- | --- |
|  |  | locally-best | design-free |
| Gene1 | 6.336E-02 | 3.119E-03 | 3.109E-03 |
| Gene2 | 3.700E-08 | 8.469E-03 | 8.193E-03 |
| Gene3 | 5.770E-05 | 7.831E-01 | 7.905E-01 |
| Gene4 | 2.465E-03 | 2.245E-03 | 2.416E-03 |

**TABLE 3.** Effect Sizes on the Illustrative Dataset

|  | log fold | Van Elteren |  |
| --- | --- | --- | --- |
|  |  | locally-best | design-free |
| Gene1 | 0.673 ↑ | 0.375 ↓ | 0.375 ↓ |
| Gene2 | -1.182 ↓ | 0.611 ↑ | 0.611 ↑ |
| Gene3 | -3.664 ↓ | 0.488 ↓ | 0.489 ↓ |
| Gene4 | -0.817 ↓ | 0.373 ↓ | 0.372 ↓ |

#### 3.1.1 Suppressing False Negatives

Batch effect may introduce false negatives, where a significantly differentially expressed gene is overshadowed. For gene 1, on which cell type B always have higher expression on both patients, the Wilcoxon rank-sum test did not pass the threshold of 0.05 to reject the null hypothesis, while Van Elteren test yields a significant p-value. The effect size, smaller than 0.5, also correctly suggests that the expression of gene 1 in cell type B is higher than that in cell type A, compared with the average logarithmic fold change, which wrongfully indicates otherwise.

#### 3.1.2 Suppressing Reversed Conclusions

Batch effect may also lead to reversed conclusion (i.e., which cell type has higher expression). For gene 2, on which cell type A always have higher expression value, both tests reject the null hypothesis. However, the effect size of Van Elteren test, larger than 0.5, correctly identifies that the expression of gene 2 in cell type A is higher than that in cell type B, while the average logarithmic fold change wrongfully indicates otherwise.

#### 3.1.3 Suppressing False Positives

False discoveries are also possible outcome of batch effect. As is shown in gene 3, the distribution of both cell types are exactly the same in each patient. Nevertheless, Wilcoxon test yields a very significant p-value. The average logarithmic fold change also has a large magnitude. Van Elteren test returns a p-value of 0.7831, together with a effect size close to 0.5, suggesting that the difference is neither significant nor large.

#### 3.1.4 Consistency

As a negative control, when the three issues above are not present, p-value from Van Eleteren test is consistent with Wilcoxon rank-sum test, as is shown by gene 4. The effect size and the log fold change also both show that the cell type B has higher expression in gene 4.

### 3.2 Simulation Study

To evaluate the performance of differential analysis methods, we simulated scRNA-seq datasets using Splatter [22]. The expression of each gene in each cell follows a Poisson–gamma mixture (i.e., negative binomial distribution), whose parameters follow a hierarchical model which characterizes the library size for each cell using a log-normal distribution, and the expression level of a gene using another gamma distribution. In addition, outlier genes are chosen by a Bernoulli distribution and scaled by a log-normal distribution, and a mean-variance trend in the expression is enforced by simulated Biological Coefficient of Variation. Using the R package of Splatter, 1,000 genes were simulated for six samples, each containing 100, 200, 300, 400, 500, and 600 cells. The split of two cell types is 3:7. Genes are randomly (*p* = 0.1) designated as differentially expressed and multiplied by a factor following a log-normal distribution (*σ* = 0.4, *μ* as specified below). For other parameters, we used default values except for `batch.facLoc=2.0`, `batch.facScale = 0.8`, and `bcv.common = 0.5`. The genes that are expected to be differentially expressed was identified using the simulated parameters and served as the ground truth for our assessment.

We compared Van Elteren test with all state-of-the-art differential expression analysis methods that interface with Seurat, i.e., Wilcoxon Rank Sum test, log likelihood test, ROC Analysis, Student’s t-test, Negative Binomial test, Poisson Test, Logistic Regression, MAST, and DESeq2. We used raw data whenever possible, as suggested by Luecken and Theis [13], except for MAST, which runs only on log-transformed normalized data. Batch information were input for Van Elteren test, Linear Regression test, Negative Binomial test, Poisson test, and MAST, which have the ability to account for batch effects.

We used the receiver operating curve (ROC) for each test to compare the methods intuitively. Formally, the ROC is the trace of false positive rate (FPR) and true positive rate (TPR) for selecting different numbers of top genes ordered by their p-values (or power for ROC Analysis). The area under curve (AUC) of the ROC was used as a quantitative metric. A good statistical test would achieve a high true positive ratio at a low false positive ratio, and thus have a greater AUC.

One example (*μ* of the log-normal distribution for DEG was set to 0.1) is shown in Fig. 3. Van Elteren test is clearly better than all methods not accounting for the batch effect (dashed lines). By a smaller margin, Van Elteren test also achieves greater AUC (0.860) than MAST (0.841), negative binomial (0.839), logistic regression test (0.833), and Poisson test (0.763), where the batch effects are accounted for as covariates. It is notable that tests uses covariates or stratification generally performs better than those that do not, showing the importance of accounting for such factors when analyzing dataset with batch effects. An exception is Poisson test, because the dataset largely deviates from the Poisson distribution. Nevertheless, Poisson test with covariates still performs better than its counterpart without covariates.

**Fig. 3.**
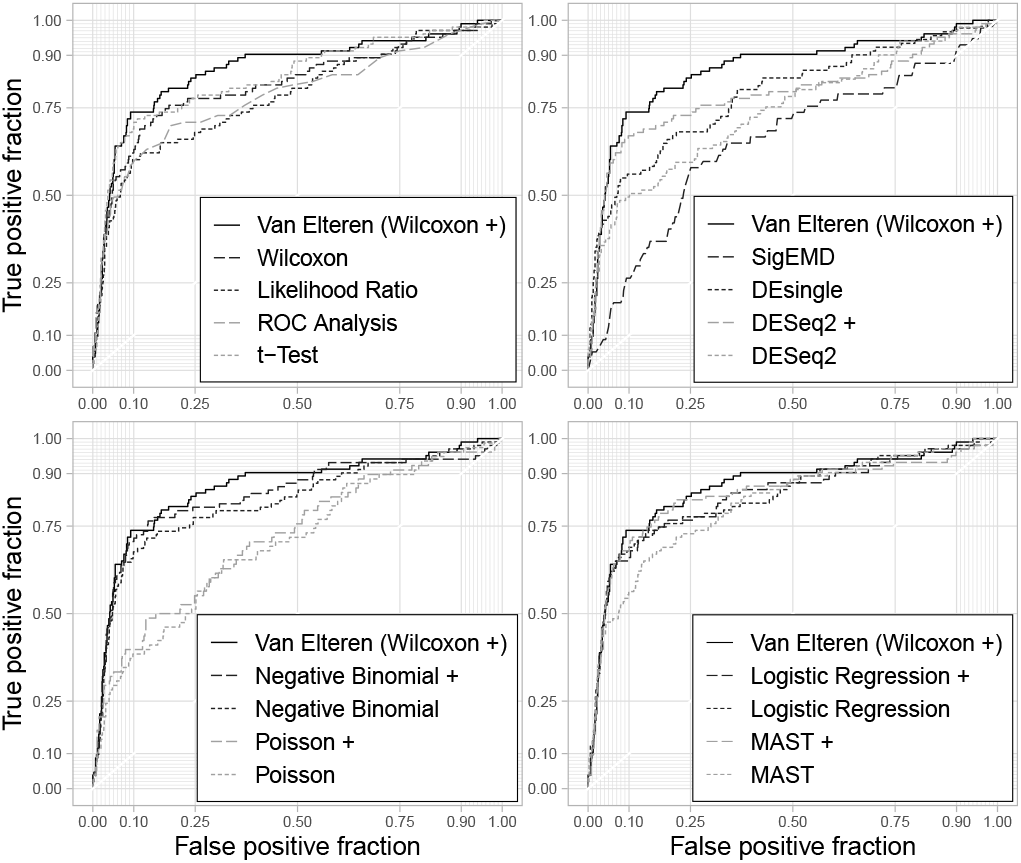
ROC curve of tested methods. Solid lines are for stratified tests or test with covariates (labeled by “+” symbol in the legends), while dashed lines are for the others. Methods are split into two panels for visual clarity.

To confirm the observation, we used *μ* = 0.1, 0.3, and 0.5 to repeat the experiment. Results summarized in Fig. 4 consolidate that tests uses covariates or stratification generally performs better. With the batch effects addressed, the accuracy of Van Elteren test, linear regression test, negative binomial test, and MAST are largely comparable. Van Elteren test performs slightly better in terms of median/mean AUC with a smaller variance. The performances of all methods improve when the effect size (which corresponds to *μ*) is larger, while the contrasts of the methods remain unchanged. It should be noted that gamma–Poisson mixture is a major part of Splatter, which naturally favors negative binomial test and MAST. In reality, patterns of gene expression are highly variable. The performance of parametric methods will decrease significantly, like the Poisson test, when the model does not fit. The nonparametric Van Elteren test is more versatile to different distributions.

**Fig. 4.**
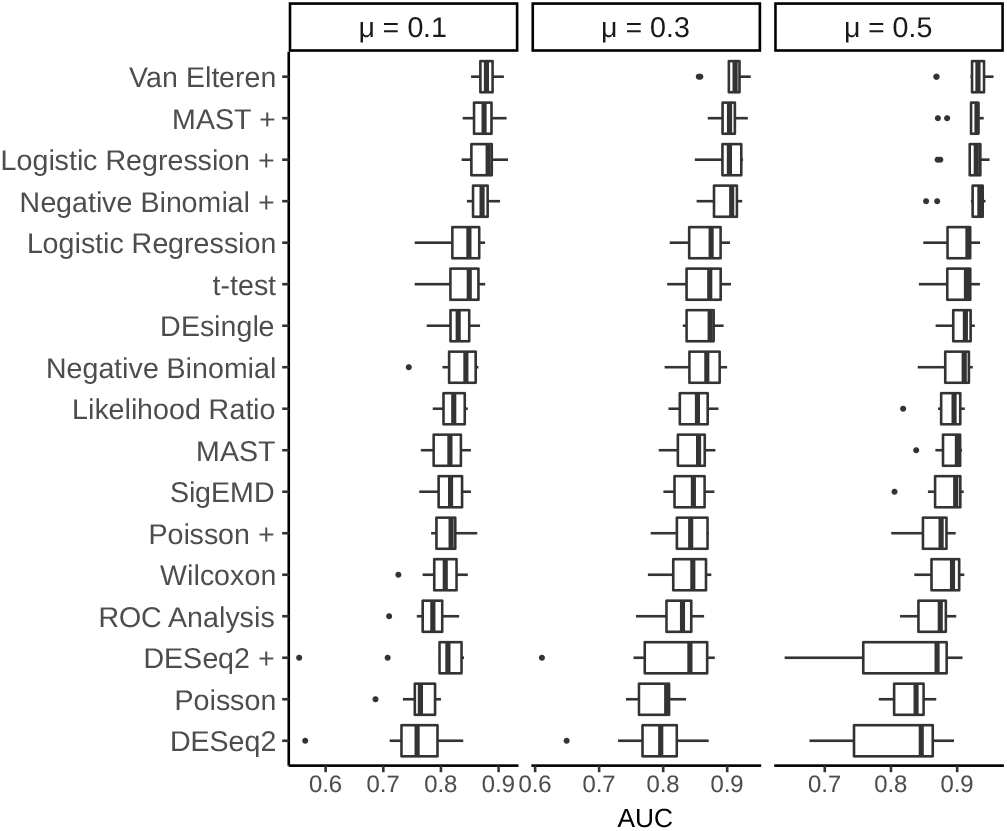
AUC of tested methods in *n* = 10 randomly simulated datasets. The median (center lines), interquartile range (hinges), 1.5 interquartile range (whiskers), and corresponding data points (dots) are shown. “*μ*” is the location parameter of the log-normal distribution for DEG.

### 3.3 Retina Data

We have tested the Van Elteren test on real retina singlecell RNA sequencing data gathered from three patients [23]. Two regions, macula (i.e., the center area) and peripheral, are labeled in the data of 5,873 cells × 10,713 genes. Cell types was annotated using unbiased clustering and marker genes in the original publication. We question which genes differentially express for the same cell type between two regions. We ran Wilcoxon rank-sum test and Van Elteren test on 2,295 rod cells and 203 cone cells. Original counts (accessed by GEO series number GSE133707) were used for both methods. We compare the results in Fig. 5, where genes with large differences in p-values between two tests are labeled with gene names.

**Fig. 5.**
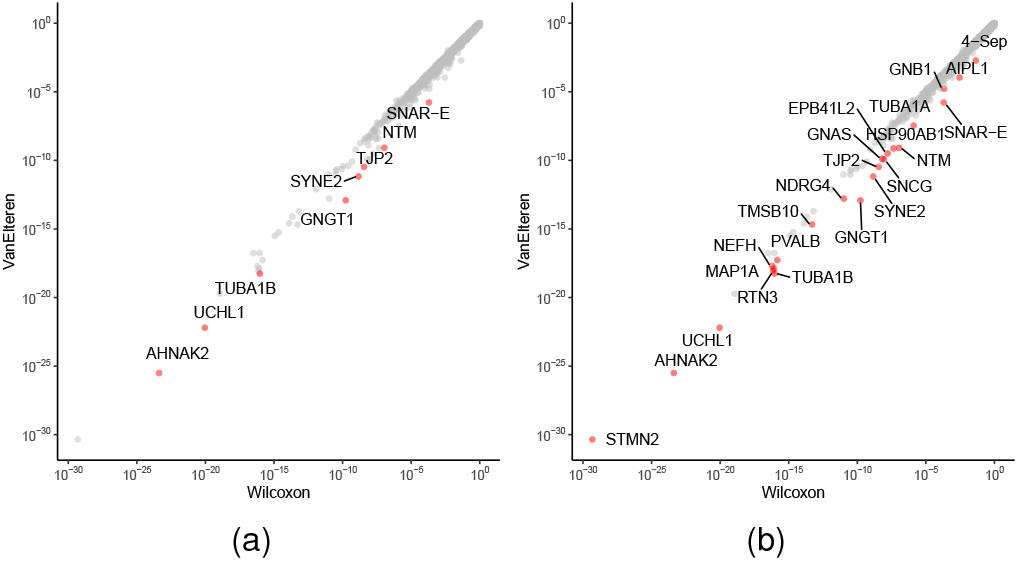
Comparison of p-values returned from Wilcoxon rank-sum test and Van Eletern test on (a) rod cells and (b) cone cells. Each dot is a gene, whose p-value from Wilcoxon rank-sum test and Van Elteren test are shown by its x-coordinate and y-coordinate, respectively. Genes with largely changed p-values (10^2^ for rod cells and 10^1^ for cone cells) are labeled with gene names.

#### 3.3.1 Rod Cells

The results of two tests are largely comparable, showing a diagonal pattern. Meanwhile, some exceptions are present (see Table 4 for the p-values and effect sizes), among which we observed that p-values for gene *GNGT1* and *SYNE2* change the most.

**TABLE 4.**
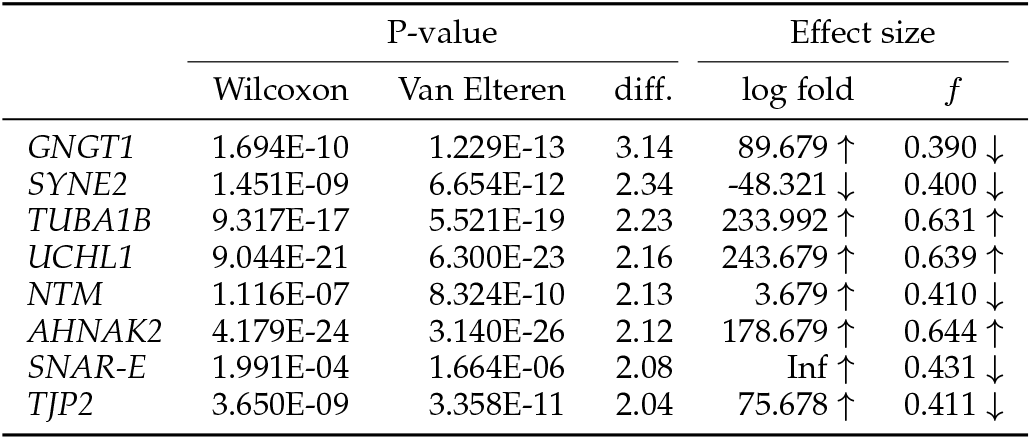
Gene with large p-value chagne in rod cells

|  | P-value |  |  | Effect size |  |
| --- | --- | --- | --- | --- | --- |
| | Wilcoxon | Van Elteren | diff. | log fold | $f$ |
| *GNGT1* | 1.694E-10 | 1.229E-13 | 3.14 | 89.679 ↑ | 0.390 ↓ |
| *SYNE2* | 1.451E-09 | 6.654E-12 | 2.34 | -48.321 ↓ | 0.400 ↓ |
| *TUBA1B* | 9.317E-17 | 5.521E-19 | 2.23 | 233.992 ↑ | 0.631 ↑ |
| *UCHL1* | 9.044E-21 | 6.300E-23 | 2.16 | 243.679 ↑ | 0.639 ↑ |
| *NTM* | 1.116E-07 | 8.324E-10 | 2.13 | 3.679 ↑ | 0.410 ↓ |
| *AHNAK2* | 4.179E-24 | 3.140E-26 | 2.12 | 178.679 ↑ | 0.644 ↑ |
| *SNAR-E* | 1.991E-04 | 1.664E-06 | 2.08 | Inf ↑ | 0.431 ↓ |
| *TJP2* | 3.650E-09 | 3.358E-11 | 2.04 | 75.678 ↑ | 0.411 ↓ |

For *GNGT1*, the reversed conclusion effect is also observed, as the Van Elteren *f* effect size suggests that the peripheral region has a higher expression, while the log fold change indicates otherwise. We further inspected the distributions to validate and interpret the differences. In Fig. 6A, the left panel does show generally higher expression of *GNGT1* in each individual patient, while the aggregated distribution on the right shows a reversed effect, which is an instance of the aforementioned Simpson’s paradox. For *SYNE2* (Fig. 6B), conspicuous discrepancy among batches is also shown, which leads to a less precise rank-sum test result. Indeed, these two genes were found playing roles in macular degeneration diseases [24], [25].

**Fig. 6.**
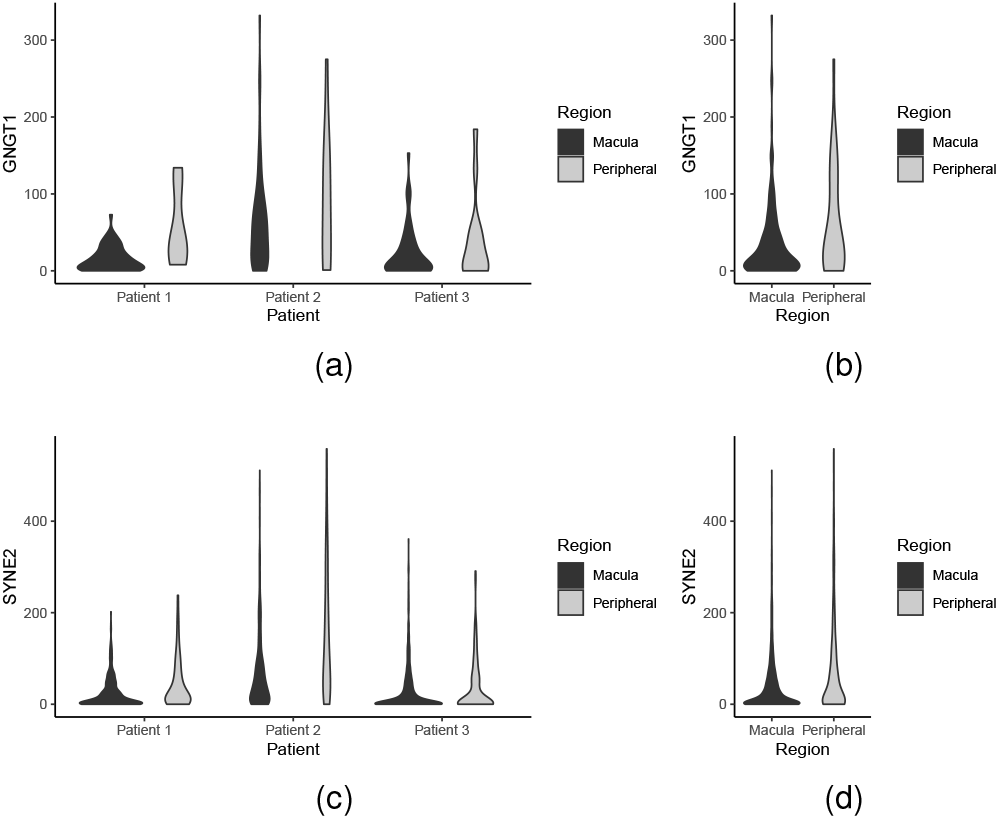
Distribution of counts of (a)(b) *GNGT1* and (c)(d) *SYNE2* in rod cells. Left panels are stratified by patients and right panels show aggregated distributions.

#### 3.3.2 Cone Cells

For cone cells, the results are similar. Some genes show changes while most genes are consistent across the tests. The p-values and effect sizes are shown in Table 5.

**TABLE 5.** Gene with large p-value change in cone cells

|  | P-value |  |  | Effect size |  |
| --- | --- | --- | --- | --- | --- |
| | Wilcoxon | Van Elteren | diff. | log fold | $f$ |
| *PCBP4* | 1.734E-02 | 1.285E-03 | 1.13 | 5.397 ↑ | 0.335 ↓ |

For gene *PCBP4*, Van Elteren test shows more significant p-value, and an effect size indicating smaller expression in macula, which is different from the log fold change. Decrease in *PCBP4* has also been linked with age-related macular degeneration [26]. Fig. 7 shows that batch effect in distribution of *PCBP4* may have misled the rank-sum test and the logarithmic fold change.

**Fig. 7.**
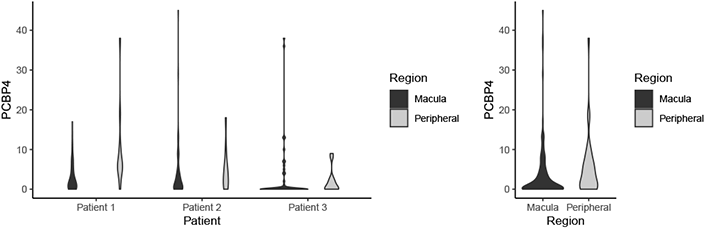
Distribution of counts of *PCBP4* in rod cells. The left panel is stratified by patients and the right panel shows aggregated distributions.

## 4 Discussion

The results have clearly shown that Van Elteren test benefits biological studies in precisely identifying differentially expressed genes. Although the results we show do not include multiple comparison correction, Seurat 3.0 will automatically give corrected p-value based on the raw p-value using Bonferroni correction. Generally, any correction based on p-values will also apply.

Our simulation study shows that Van Elteren test is one of the best performing methods among the most widely used state-of-the-art methods. It should be noted that due to no free lunch theorem, the performance of a method depends on the degree of match of the data and the model. Being a nonparametric test, Van Elteren test does not assume a specific distribution and thus suits the exploratory analysis of a dataset, where the distribution of the gene expression varies. This is especially valuable for scRNA-seq data analyses as it is cumbersome to check the distribution pattern of thousands of genes.

The result also indicates that stratified test is a neat way to handle batch effect. Although covariate has the ability to control for explanatory variables, it is generally more suitable for continuous variables. It also casts more assumptions when modeling covariate. Stratified test, on the other hand, does not infer the influence of the discrete batches. Rather, it directly aggregates the statistical power of multiple samples.

We also show that the weighted common language effect size, a byproduct of Van Elteren test, reflects the difference of gene expression more faithfully. It can be used with Van Elteren test or other tests, to support a more comprehensive understanding of the data.

Admittedly, for the rod cells in the retinal data, although changes in p-values are observed, the significance threshold was well passed by both. However, it should be noted that the retina data are collected from relatively healthy tissues and are considered clean, while Van Elteren test is expected to make a more meaningful difference on noisy pathological and tumor data. In addition, rod cell is the most populous cell in retina. For rare cell types that take smaller proportions, like the cone cells, the difference Van Elteren test makes can be crucial.

Overall, Van Elteren test and our modified common language effect size are direct extensions of the Wilcoxon ranksum test and common language effect size. As nonparametric methods, they perform well in various scenarios, with or without obvious batch effect. A caveat of the stratified test is that for it to work the strata shall not overlap with the variable of interest. For instance, it may not find the difference, meanwhile also control for the batch effect, between two patients. Nonetheless, neither is covariate applicable to such cases. As the batch effect and biological effect are convoluted, more prior knowledge is generally needed to distinguish them. Another limitation is that it does not detect interaction effects between variables, which may need more complex nonparametric tests such as Aligned Rank Transform [27]. Besides, when the distribution can be reasonably assumed, such as a normal distribution where the central limit theorem apply, a corresponding parametric test may offer a higher power.

## 5 Summary

We have adopted Van Elteren test, an underappreciated statistical test, and our weighted common language effect size to single-cell sequencing data. When batch effect is severe, the test control for false positives and false negatives. Otherwise, it is consistent with Wilcoxon rank-sum test. Simulation study show that Van Elteren test achieves the state of the art. The modified common language effect size also faithfully depicts the trends. This work may increase the precision of differential expression analysis to help identify genes of interests.

## Acknowledgments

This work was supported in part by grant number 2018-182735 to KC, Human Breast Cell Atlas Seed Network Grant (HCA3-0000000147) to KC from the Chan Zuckerberg Initiative DAF, an advised fund of Silicon Valley Community Foundation, grant RP180248 to KC from Cancer Prevention & Research Institute of Texas, The University of Texas MD Anderson Cancer Center Pre-Cancer Atlas Project to KC, and The University of Texas MD Anderson Cancer Center Colorectal Cancer Moonshot and P30 CA016672 (US National Institutes of Health/National Cancer Institute) to the University of Texas Anderson Cancer Center Core Support Grant.

## References

[1] A. Regev, S. A. Teichmann, E. S. Lander, I. Amit, C. Benoist, E. Birney, B. Bodenmiller, P. Campbell, P. Carninci, M. Clatworthy et al., “Science Forum: the Human Cell Atlas,” Elife, vol. 6, p. e27041, 2017.

[2] C. R. Blyth, “On simpson’s paradox and the sure-thing principle,” Journal of the American Statistical Association, vol. 67, no. 338, pp. 364–366, 1972.

[3] F. Wilcoxon, “Individual comparisons by ranking methods,” in Breakthroughs in statistics. Springer, 1992, pp. 196–202.

[4] H. B. Mann and D. R. Whitney, “On a test of whether one of two random variables is stochastically larger than the other,” The annals of mathematical statistics, pp. 50–60, 1947.

[5] P. Van Elteren, “On the combination of independent two-sample tests of Wilcoxon,” Bull Inst Intern Staist, vol. 37, pp. 351–361, 1960.

[6] D. S. Kerby, “The simple difference formula: An approach to teaching nonparametric correlation,” Comprehensive Psychology, vol. 3, pp. 11–IT, 2014.

[7] K. O. McGraw and S. Wong, “A common language effect size statistic.” Psychological bulletin, vol. 111, no. 2, p. 361, 1992.

[8] L. Haghverdi, A. T. Lun, M. D. Morgan, and J. C. Marioni, “Batch effects in single-cell RNA-sequencing data are corrected by matching mutual nearest neighbors,” Nature biotechnology, vol. 36, no. 5, p. 421, 2018.

[9] B. Hie, B. Bryson, and B. Berger, “Efficient integration of heterogeneous single-cell transcriptomes using Scanorama,” Nature biotechnology, vol. 37, no. 6, p. 685, 2019.

[10] T. Stuart, A. Butler, P. Hoffman, C. Hafemeister, E. Papalexi, W. M. Mauck III, Y. Hao, M. Stoeckius, P. Smibert, and R. Satija, “Comprehensive integration of single-cell data,” Cell, 2019.

[11] I. Korsunsky, N. Millard, J. Fan, K. Slowikowski, F. zhang, K. Wei, Y. Baglaenko, M. Brenner, P.-r. Loh, and S. Raychaudhuri, “Fast, sensitive and accurate integration of single-cell data with Harmony,” Nature methods, pp. 1–8, 2019.

[12] V. Nygaard, E. A. Rødland, and E. Hovig, “Methods that remove batch effects while retaining group differences may lead to exaggerated confidence in downstream analyses,” Biostatistics, vol. 17, no. 1, pp. 29–39, 2016.

[13] M. D. Luecken and F. J. Theis, “Current best practices in singlecell rna-seq analysis: a tutorial,” Molecular systems biology, vol. 15, no. 6, p. e8746, 2019.

[14] A. McDavid, G. Finak, P. K. Chattopadyay, M. Dominguez, L. Lamoreaux, S. S. Ma, M. Roederer, and R. Gottardo, “Data exploration, quality control and testing in single-cell qpcr-based gene expression experiments,” Bioinformatics, vol. 29, no. 4, pp. 461–467, 2013.

[15] G. Finak, A. McDavid, M. Yajima, J. Deng, V. Gersuk, A. K. Shalek, C. K. Slichter, H. W. Miller, M. J. McElrath, M. Prlic et al., “Mast: a flexible statistical framework for assessing transcriptional changes and characterizing heterogeneity in single-cell rna sequencing data,” Genome biology, vol. 16, no. 1, pp. 1–13, 2015.

[16] M. I. Love, W. Huber, and S. Anders, “Moderated estimation of fold change and dispersion for rna-seq data with deseq2,” Genome biology, vol. 15, no. 12, p. 550, 2014.

[17] V. Svensson, “Droplet scrna-seq is not zero-inflated,” Nature Biotechnology, vol. 38, no. 2, pp. 147–150, 2020.

[18] B. Everitt and A. Skrondal, The Cambridge Dictionary of Statistics, Fourth Edition, ser. BusinessPro collection. Cambridge University Press, 2010.

[19] R. Lopez, J. Regier, M. B. Cole, M. I. Jordan, and N. Yosef, “Deep generative modeling for single-cell transcriptomics,” Nature methods, vol. 15, no. 12, p. 1053, 2018.

[20] T. Wang and S. Nabavi, “Sigemd: A powerful method for differential gene expression analysis in single-cell rna sequencing data,” Methods, vol. 145, pp. 25–32, 2018.

[21] Z. Miao, K. Deng, X. Wang, and X. Zhang, “Desingle for detecting three types of differential expression in single-cell rna-seq data,” Bioinformatics, vol. 34, no. 18, pp. 3223–3224, 2018.

[22] L. Zappia, B. Phipson, and A. Oshlack, “Splatter: simulation of single-cell rna sequencing data,” Genome biology, vol. 18, no. 1, pp. 1–15, 2017.

[23] Q. Liang, R. Dharmat, L. Owen, A. Shakoor, Y. Li, S. Kim, A. Vitale, I. Kim, D. Morgan, S. Liang, N. Wu, K. Chen, M. M. DeAngelis, and R. Chen, “Single-nuclei RNA-seq on human retinal tissue provides improved transcriptome profiling,” Nature Communications,vol. 10, no. 1, pp. 1–12, 2019.

[24] A. V. Kolesnikov, L. Rikimaru, A. K. Hennig, P. D. Lukasiewicz, S. J. Fliesler, V. I. Govardovskii, V. J. Kefalov, and O. G. Kisselev, “G-protein *βγ*-complex is crucial for efficient signal amplification in vision,” Journal of Neuroscience, vol. 31, no. 22, pp. 8067–8077, 2011.

[25] D. M. Maddox, G. B. Collin, A. Ikeda, C. H. Pratt, S. Ikeda, B. A. Johnson, R. E. Hurd, L. S. Shopland, J. K. Naggert, B. Chang et al., “A mutation in *Syne2* causes early retinal defects in photoreceptors, secondary neurons, and müller glia,” Investigative ophthalmology & visual science, vol. 56, no. 6, pp. 3776–3787, 2015.

[26] J. G. Meyer, T. Y. Garcia, B. Schilling, B. W. Gibson, and D. A. Lamba, “Proteome and secretome dynamics of human retinal pigment epithelium in response to reactive oxygen species,” Scientific reports, vol. 9, no. 1, pp. 1–12, 2019.

[27] J. O. Wobbrock, L. Findlater, D. Gergle, and J. J. Higgins, “The aligned rank transform for nonparametric factorial analyses using only anova procedures,” in Proceedings of the SIGCHI conference on human factors in computing systems, 2011, pp. 143–146.

